# MITF regulates gene expression in middle tufted neurons and other projection neurons of the olfactory bulb

**DOI:** 10.1101/2025.01.29.635553

**Authors:** Fatich Mechmet, Alba Sabaté San José, Snævar Sigurðsson, Ingvi Gautsson, Eiríkur Steingrímsson, Pétur Henry Petersen

**Affiliations:** Department of Anatomy, BioMedical Center, Faculty of Medicine, University of Iceland, Reykjavik, Iceland; School of Health Sciences, BioMedical Center, University of Iceland, Reykjavik, Iceland; Utopia Arctica, Reykjavik, Iceland; Department of Biochemistry and Molecular Biology, BioMedical Center, Faculty of Medicine, University of Iceland, Reykjavik, Iceland

**Keywords:** MITF, olfactory bulb, projection neurons, middle tufted cells, neuronal activity

## Abstract

The microphthalmia-associated transcription factor (MITF) is a master transcription factor in melanocytes and plays equally important roles in mast cells. It is required for the generation, differentiation, and function of these cell types in vertebrates. *Mitf* is also expressed in the projection neurons of the olfactory bulb (OB), the mitral and tufted (M/T) cells. Loss of *Mitf* leads to neuronal hyperactivity in primary M/T cells but the general function of *Mitf* in neurons is unknown. Here, we identify putative MITF target genes in M/T cells, which show limited overlap with known targets in other cell types. These genes can be divided into two groups, those likely to inhibit neuronal activity and genes expressed specifically in a subclass of tufted cells, the middle tufted cells (mTCs). The mTCs are reduced in number in the *Mitf* mutant OB, suggesting a role for *Mitf* in the generation or survival of the mTCs and/or their function.

**Significance statement:** Identifying MITF target genes in the OB offers insights into its role in olfaction and the regulation of neuronal activity as well as uncovering its role in mTCs. MITF may play a role in various other processes in neurons.

## Introduction

A single gene can play a central role in multiple cellular and developmental contexts. One such gene is the microphthalmia-associated transcription factor (*Mitf*), which is often referred to as the master regulator of melanocytes as it is required for their generation and function (Goding & Arnheiter, 2019). *Mitf* is also a central gene in retinal pigment epithelial cells (RPE), mast cells and osteoclasts (Goding & Arnheiter, 2019; Steingrímsson et al., 2004). Mice homozygous for the *Mitf^mi-vga9^* null mutation are white due to lack of melanocytes (Hodgkinson et al., 1993), they lack mast cells (Ingason et al, 2019), have small eyes due to RPE defects, their osteoclasts are immature and they show hearing impairments due to absence of melanocytes in the inner ear (Holtrop et al., 1981; Morii et al., 2004; Steingrímsson et al., 2004). *Mitf* is also a well-characterized melanoma oncogene (Goding & Arnheiter, 2019). The MITF protein is a transcription factor and, as such, has several cell-type specific target genes. These include *Dct*, *Pmel*, *Mlana*, *Trpm1*, and *Tyrp1* in melanocytes (Goding & Arnheiter, 2019) and several mast-cell specific genes including proteases (tryptases, chymases), adhesion molecules (integrin α4 subunit, SGIGSF, PAI-1), metabolic enzymes (*granzyme B*, TPH, Hdc, PDG2 synthase) and growth factor receptors (*NGFR*, *Kit*) (Oppezzo & Rosselli, 2021). Recently, *Mitf* was shown to play a central role in nerve repair in the peripheral nervous system (Daboussi et al., 2023). *Mitf* is also expressed in the mammalian central nervous system (CNS), distinctly in mitral and tufted (M/T) cells of the olfactory bulb (OB) (Ohba et al., 2015; Atacho et al. 2020). These cells are the projection neurons (PNs) of the OB, which link the OB to cortical areas and transduce olfactory information. Loss of *Mitf* does not affect the overall architecture of the OB but leads to hyperactivity in primary M/T cells, likely due to a decrease in expression of *Kcnd3,* the gene coding for the potassium channel subunit Kv4.3 (Atacho et al., 2020). Apart from *Kcnd3,* MITF target genes in neurons are unknown. Importantly, besides its involvement in neuronal activity, MITF likely plays a role in other cellular processes of M/T cells, similar to its diverse functions in melanocytes and other cell types. The aim of the current study is to further identify putative MITF target genes in the PNs of mouse OB.

## Material and Methods

### Animals

All *in vivo* procedures were approved by the Committee on Experimental Animals and were in accordance with the Regulation 460/2017 and European Union Directive 2010/63 (license number: 2013-03-01). Wild type (C57BL/6J), and homozygous *Mitf^mi-vga9^* mutant mice were used in this study. The *Mitf^mi-vga9^*mutation is a loss of function mutation caused by transgene insertion that affects the expression of MITF (Hodgkinson et al., 1993; Tachibana et al., 1992). The animals were kept in the mouse facility of the University of Iceland. The animals were housed in groups of 2-4 per cage under controlled conditions (21-22°C; 12 h light/ 12 h dark), and food and water provided *ad libitum* unless otherwise specified. Mice of both genders were used for odor exposure, RNA sequencing and single-molecule fluorescence *in situ* hybridization (smFISH) experiments.

### Odor exposure

The odor exposure experiment was performed in two experimental setups. In both setups, mice aged 2-3 months old were kept in cages in an odor-free room. An Eppendorf tube containing 120 μL of amyl-acetate (AA); Sigma-Aldrich, cat# W504009) was placed in the cage for 30 minutes at nose level. In the first experimental setup, mice were immediately sacrificed after being exposed to AA for 30 minutes. In the second experimental setup, the tube was removed from the cage and mice were kept in an odor-free room for an additional 30 minutes before being sacrificed. The OBs were collected and flash-frozen in liquid nitrogen for total RNA extraction.

### Cell culture

N2a cells were purchased from the Leibniz Institute DSMZ-Deutsche Sammlung von Mikroorganism und Zellkulturen GmbH. The cells were cultured in Dulbeccós Modified Eagle Medium (DMEM)-GlutaMAX (Gibco, cat# 31966-021) supplemented with 10% Fetal Bovine Serum (FBS; #10270-106, Gibco) and 1% MEM Non-Essential Amino Acids solution (NEAA; Gibco, cat# 11140-050). The cells were grown in 75 cm^2^ Nunc EasYFlasks with Nunclon Delta surface culture dishes (Thermo Scientific, cat# 156499) at 37°C with 5% CO_2_. The cells were passaged at a 1:9 dilution after they reached 70-80% confluency by gentle trypsinization.

### Generation of stable doxycycline inducible N2a cells overexpressing MITF

N2a cells were grown in 75 cm^2^ Nunc EasYFlask in Nunclon Delta surface culture dishes (Thermo Scientific, cat# 156499) to 70-80% confluency and transfected with a mixture of three piggy-Bac vectors (0.75 μg transposase [K7.7 pBase], 0.075 μg reverse tetracycline transcription construct activator [rtTA], and 0.75 μg PB-CMW-MITF-flag-HA [6430 bp]) using FuGENE HD transfection reagent (Promega, cat# E2311). Cells were transfected with K7.7 pBase, rtTA and PB-EV-3XFLAG-HA (5152 bp) was used as control (Möller et al., 2019). Following 48 hours of transfection, the cells were treated with 0.55 mg/mL Geneticin (G418) (Gibco, #10131-035) to select successfully transfected cells.

### Western blotting

N2a cells were seeded in Nunclon 6-well plates (Thermo Scientific, cat# 140675) at 3.5 x 10^5^ per well overnight (O/N). After inducing differentiation by O/N serum deprivation, the cells were treated with 0.5 µg/mL doxycycline (dox; SCBT, cat# sc-204734B) O/N. The cells were then harvested with 50 µL/well radioimmunoprecipitation assay (RIPA) buffer (250 mM NaCl, 1% IGEPAL CA-630, 0.5% sodium deoxycholate, 0.1% sodium dodecyl sulphate, 50 mM Tris-HCl pH 8.0) containing 1x Halt protease and phosphatase inhibitor cocktail (Thermo Scientific, #78440). After centrifugation at 16,000 x g for 20 minutes at 4°C, samples were diluted with 2x sample buffer (4% sodium dodecyl sulphate, 20% glycerol, 0.02% bromophenol blue, 120 mM Tris-HCl pH 6.8) containing 5% β-mercaptoethanol (Aldrich, cat# M6250) at 1:1 ratio and lysates were heated at 95°C for 5 minutes. The protein lysates were run on 12.5% SDS-PAGE gel and transferred onto 0.2 µm PVDF membrane (Thermo Scientific, cat# 88520). The membrane was incubated in a blocking buffer consisting of 5% BSA (Carl Roth, cat# 3737.4) in 1x Tris-buffered saline/0.1% Tween 20 (TBS-T) for 1 hour at room temperature (RT). Following blocking, the membranes were incubated O/N at 4°C with the following primary antibodies diluted in blocking solution: mouse monoclonal IgG1 anti-FLAG M2 (1:2,500; Sigma-Aldrich, cat# F1804) and rabbit polyclonal IgG anti-β-Tubulin (1:5,000; abcam, cat# ab15568). Membranes were washed with 1x TBS-T and incubated for 1 hour at RT with the following secondary antibodies diluted at 1:10,000 in blocking buffer: IRDye goat anti-mouse IgG1 800 (LI-COR Biosciences) and IRDye donkey anti-rabbit IgG 680 (LI-COR Biosciences). After washing steps with 1x TBS-T, the membranes were imaged using Odyssey CLx Imager (LI-COR Biosciences).

### RNA sequencing (RNA-seq) and gene expression analysis

For RNA sequencing, RNA was extracted from OB tissues of 1-2 months old C57BL/6J and *Mitf^mi-vga9/mi-vga9^*mice and from the N2a cell line using Monarch Total RNA Miniprep Kit (New England Biolabs Inc., cat# T2010S). RNA integrity (RIN) of the total RNA was determined using the DNA 5K/RNA chip on the LabChip GX instrument. TruSeq RNA v2 Sample Prep Kit (Illumina, RS-122-2001) was used to generate cDNA libraries derived from Poly-A mRNA isolated from total RNA samples (0.2-1 μg) and hybridized to Poly-T beads. After the Poly-A mRNA fragmentation at 94°C in the presence of divalent cations, first-strand cDNA synthesis was performed using random hexamers and SuperScript IV reverse transcriptase (Invitrogen, cat# 18090010). Following second-strand cDNA synthesis and PCR amplification, cDNA sequencing libraries were created. The libraries were measured on the LabChip GX. Following this, they were diluted to 3 nM and stored at-20°C until use. Samples were pooled and clustered on NovaSeq S4 flow cells (Illumina). Paired-end sequencing was performed using XP workflow on NovaSeq 6000 instruments (Illumina). Base-calling was performed in real-time using RTA v3.4.4 demultiplexing of BCL file and bcl2fastq2 v2.20 was used to generate FASTQ files.

For transcript level analysis, paired-end RNA-seq reads were pseudoaligned to the mouse reference genome (*Mus musculus*. GRCm38.96) (Yates et al., 2016) using a companion package of Kallisto (v 0.46.1) (Bray et al., 2016). Kallisto output was analyzed in Sleuth (Pimentel et al., 2017) to determine differentially expressed (DE) genes in the OBs of C57BL/6J and *Mitf^mi-vga9/mi-vga9^*mice (n=6 per genotype), between N2a (MITF) (n=3) and N2a (EV) (n=3). In the two odor exposure experiments (30 minutes and 60 minutes odor exposure), three untreated C57BL/6J and *Mitf^mi-vga9/mi-vga9^* mice were used as controls and three mice per genotype were exposed to odor (AA) in each odor exposure experiment. The significance of DE genes (p-/q-value) and the fold change (beta estimate) was determined using likelihood test (LRT) intersected in Sleuth. After normalization of odor-exposed OBs to control OBs, odor exposure experiments were analyzed using the spline version 4.1.2 package in R to create a model for the time-course. This model was used to create the design-matrix for the Sleuth analysis that allowed testing the effect of the length of two different odor exposure experiments at the same time.

Volcano plot was generated by plotting significance on the y-axis and cut-off of fold change on the x-axis using the R packages “ggplot2” and “ggrepel”. Significance and fold change values of DE genes from RNA-seq data were as follows: q-value ≤ 0.05; cut-off of |b-value (foldchange)|≥ 0.7 for *Mitf^mi-vga9/mi-vga9^* compared to C57BL/6J; p-value ≤ 0.05; cut-off of |b-value (foldchange)|≥ 2 for *Mitf^mi-vga9/mi-vga9^* versus C57BL/6J after 30 and 60 minutes odor exposure; q-value ≤ 0.05; cut-off of |b-value (foldchange)|≥ 2 for N2a (MITF) compared to N2a (EV). Bar graphs showing TPM value of each gene between *Mitf^mi-vga9/mi-vga9^* and C57BL/6J in odor and non-odor experiments were generated using the R package “cowplot”. Dot plots to view the average expression values of signature genes in subgroups of PNs were obtained from publicly available data (Zeppilli et al., 2021).

### Single molecule fluorescence *in situ* hybridization (smFISH)

SmFISH was used to characterize the expression of genes in mouse OBs. The C57BL/6J and *Mitf^mi-vga9/mi-vga9^*mice, aged 1-2 months old, were sacrificed by cervical dislocation and OBs dissected and placed in O.C.T. compound (Tissue-Tek Sakura, #4583) in 15 x 15 x 15 mm Cryomolds (Tissue-Tek Sakura) and immediately flash-frozen in liquid nitrogen. The OBs were sectioned using Leica CM1950 cryostat at a thickness of 20 μm. Two sections per slide were placed onto 76 x 26 mm microscope slide adhesive++ (DWK Life Sciences, cat# 235502605). The samples were pre-treated according to the manufacturer’s instructions (Sample Preparation and Pre-treatment Guide for Fresh Frozen Tissue PART 1; Advanced Cell Diagnostics, cat# 320513). The smFISH was performed according to the “RNAscope Fluorescent Multiplex Kit User Manual PART 2” (Advanced Cell Diagnostics, cat# 320293) using probes *Mitf* (C1; cat# 422501; C2; cat# 422501), *Tbr2/Eomes* (C2; cat# 429641; C3; cat# 429641), *Htr5a* (C1; cat# 411151), *Vdr* (C1; cat# 524511), *Doc2g* (C3; cat# 517981), *Wif1* (C1; cat# 412361), *Tspan10* (C2; cat# 1190641), *Sgcg* (C3; cat# 488051), and *Fam84b* (C1; cat# 500991) (Table 1). Images were acquired at 30x magnification using 10 confocal Z-stacks as follows: two images, one from the lateral OB and one from the medial OB of each bulb for *Htr5a, Vdr, Doc2g, Wif1* and four images, two from the lateral and two from the medial OB, for *Tspan10, Sgcg* and *Fam84b*.

**Table 1.** The target probes used in smFISH.

| **Gene symbol** | **Species** | **Cat No** | **Entrez**<br>**Gene ID** | **Accession No** | **Target**<br>**region (bp)** |
| --- | --- | --- | --- | --- | --- |
| *Mitf* | Mouse | 422501 | 17342 | NM_00111319.1 | 821 – 1869 |
| *Tbr2/Eomes* | Mouse | 429641 | 13813 | NM_010136.3 | 1288 – 2370 |
| *Htr5a* | Mouse | 411151 | 15563 | NM_008314.2 | 169 – 1070 |
| *Vdr* | Mouse | 524511 | 22337 | NM_009504.4 | 427 – 1606 |
| *Doc2g* | Mouse | 517981 | 60425 | NM_021791.3 | 167 – 1154 |
| *Wif1* | Mouse | 412361 | 24117 | NM_011915.2 | 380 – 1394 |
| *Tspan10* | Mouse | 1190641 | 208634 | NM_145363.2 | 212 – 1291 |
| *Sgcg* | Mouse | 488051 | 24053 | NM_011892.3 | 2 – 1013 |
| *Fam84b* | Mouse | 500991 | 399603 | NM_001162926.1 | 884 – 1996 |

### Quantification of mRNA transcripts

Two strategies were used to count mRNA transcripts from smFISH images. In the first strategy, the ImageJ software was used to count *Htr5a, Vdr, Wif1*, and *Doc2g* transcripts according to the protocol described by Atacho and associates (Atacho et al., 2020). The second strategy used machine learning (ML) to count *Sgcg, Tspan10*, and *Fam84b* transcripts. The latter process involved annotating OB layers, quantifying smFISH signals, and assessing them across layers. A human expert initially annotated images using Krita and a graphics tablet to train the ML model (DeepLabv3+) for layer recognition and to manually correct errors. Data augmentation methods (image rotation, horizontal and vertical mirroring, and brightness adjustment) were applied to the training data to increase the model’s robustness, saving checkpoints at regular intervals for model selection post-training. Quantification was done with a dot counting approach in Python using the “findmaxima2d” package, an equivalent protocol to the “Find Maxima” operation in ImageJ. Image data processing included file conversion and specific ML methods for local maxima detection and dot counting.

### Quantification of *Fam84b* clusters

*Fam84b* positive cells were quantified using ML. The model was trained with various offsets and flip combinations to mitigate artifacts. Outcomes were merged using a weighted average. OpenCV functions extracted final nucleus region coordinates and saved as JSON files for analysis*. Fam84b* expression was quantified by pixel percentage above a threshold normalized by nuclear region area. Positive nuclei in OB layers were counted based on size and pixel count criteria.

## Statistical analysis

For all *in vivo* studies, a minimum of three mice per genotype were included. Quantitative results were analyzed using two-way ANOVA, using the R statistical package. To obtain p-values for ANOVA tests, multiple comparisons were carried out with Sidak’s multiple comparison. Results are represented as the mean with standard error (SE). Sample size was not pre-determined.

## Results

### Two groups of MITF target genes in OB neurons

The OB is divided into tissue layers, characterized by morphologically and functionally distinct neurons. Previous work has shown that *Mitf* is expressed in the mouse OB but is near absent from the OBs of *Mitf^mi-vga9/mi-vga9^*mice (Ohba et al., 2015; Atacho et al., 2020). SmFISH targeting all known *Mitf* isoforms shows *Mitf* co-expression with the OB PNs marker *Tbr2/Eomes* in large cells beneath the GL and in the MCL of both lateral and medial OBs from C57BL/6J mice (Figure 1A).

**Figure 1.**
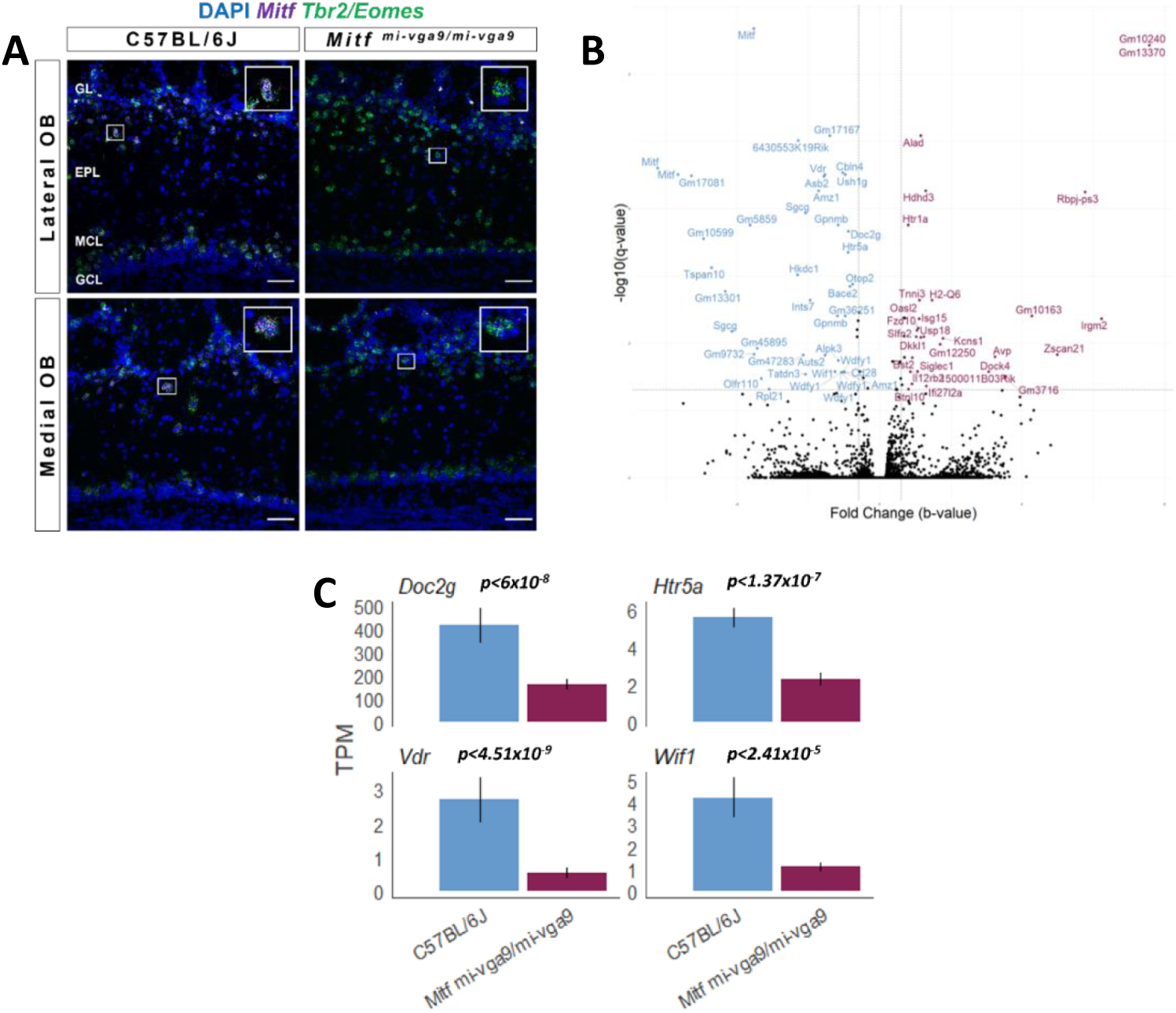
The expression of genes associated with neuronal activity is decreased in *Mitf^mi-vga9/mi-vga9^* OBs. A. Representative smFISH images from the lateral and medial OBs of C57BL/6J and *Mitf^mi-vga9/mi-vga9^* mice showing the expression of *Mitf* in *Tbr2/Eomes* positive cells. Scale bars indicate 50 μm. N=4 per genotype. B. Volcano plot shows differentially expressed genes between OBs of *Mitf^mi-vga9/mi-vga9^* and C57BL/6J mice. N=6 per genotype. C. Expression of selected genes in TPM in C57BL/6J and *Mitf^mi-vga9/mi-vga9^* OBs. N=6 per genotype. In the volcano plot, blue represents decreased genes, and purple represents increased genes. P-values (*p<0.05) on the bar graphs indicate the significance in *Mitf^mi-vga9/mi-vga9^* compared to C57BL/6J. GL (glomerular layer), EPL (external plexiform layer), MCL (mitral cell layer), GCL (granule cell layer); TPM (transcript per million).

Considering the pivotal role of the OB PNs in processing and transmitting olfactory signals, this expression pattern previously led to the hypothesis that *Mitf* might play an important role in olfaction (Atacho et al., 2020; Ohba et al., 2015). As the MITF protein regulates expression of other genes, identifying MITF target genes is the first step in understanding its function. To achieve this, comparative RNA-seq analysis was performed using OB from wild type (C57BL/6J) and *Mitf^mi-vga9/mi-vga9^* mice. Genes positively regulated by MITF should show reduced expression in the OBs of *Mitf^mi-vga9/mi-vga9^*mice, whereas suppressed genes should display increased expression. Here, the focus is on genes showing reduced expression in the absence of *Mitf*. The RNA-seq data revealed several such genes (Figure 1B, Table 2). Cross-referencing with expression information from biological and bioinformatic databases (e.g., Allen Brain Atlas, UniProt, Mouse Genome Informatics, Protein Atlas) provided a shorter list of genes with distinct and specific expression in M/T cells in C57BL/6J mice, namely *Htr5a*, *Doc2g*, *Vdr*, *Wif1, Sgcg, Tspan10* and *Olfr110*. Of these genes *Tspan10* is the only gene previously identified as a possible MITF target gene (Rambow et al. 2015). These genes can be further divided into two groups by their expression pattern – the first group is expressed exclusively in PNs in the EPL, beneath the GL (*Olfr110, Sgcg, Tspan10*), while the second group showed expression both beneath the GL and in the MCL (*Htr5a, Wif1, Doc2g, Vdr*). The global expression of the latter group in both C57BL/6J and *Mitf^mi-vga9/mi-vga9^* OBs is shown in Figure 1C. A smFISH analysis confirmed the decreased expression of *Htr5a* (Figure 2A) and *Wif1* (Figure 2B) in *Mitf^mi-vga9/mi-vga9^* OBs when compared to C57BL/6J OBs. In the OBs of *Mitf^mi-vag9/mi-vga9^*, *Htr5a* expression decreased by 73% beneath the GL and 67% in the MCL. *Wif1* expression was reduced by 89% beneath the GL and 84% in the MCL in *Mitf^m-ivga9/mi-vga9^* compared to C57BL/6J. Statistical analysis showed significant differences both in *Htr5a and Wif1* expression beneath the GL and in the MCL of *Mitf^mi-vga9/mi-vga9^* OBs (Figure 2C; *Htr5a:* beneath GL: *t _(8)_ = 6.894*, *p = 0.0008;* MCL: *t _(8)_ = 8.300*, *p = 0.0002*) (Figure 2D; *Wif1:* beneath GL: *t _(8)_ = 3.882*, *p = 0.0277*; MCL: *t _(8)_ = 9.584*, *p = 0.0001*). The smFISH analysis also showed a decrease in the expression of *Vdr* (Figure 3A) and *Doc2g* (Figure 3B). The expression of *Vdr* was reduced by 68% beneath the GL and in the MCL and statistical analysis confirmed significant differences in both layers in *Mitf^mi-vga9/mi-vga9^* compared to C57BL/6J OBs (Figure 3C; *Vdr:* beneath GL: *t _(8)_ = 8.189*, *p = 0.0002*; MCL: *t _(8)_ = 8.816*, *p = 0.0001*). Although *Doc2g* expression was reduced by 73% beneath the GL and 66% in the MCL in *Mitf^mi-vga9/mi-vga9^*OBs, statistical analysis showed no significant differences either beneath the GL or in the MCL (Figure 3D; *Doc2g:* beneath GL: *t _(8)_ = 3.070*, *p = 0.0886*; MCL: *t _(8)_ = 3.350*, *p = 0.0590*). Analysis of ChIP-seq and CUT&RUN data from melanoma cells show that *Htr5a*, *Sgcg* and *Tspan10* have MITF binding peaks in these cells (Supplementary figure 1A-C; Dilshat et al., 2021; Laurette et al., 2015; Strub et al., 2011). However, this is not the case for all the potential target genes in OBs. For instance, no peaks were identified in the *Doc2g* and *Vdr* genes.

**Figure 2.**
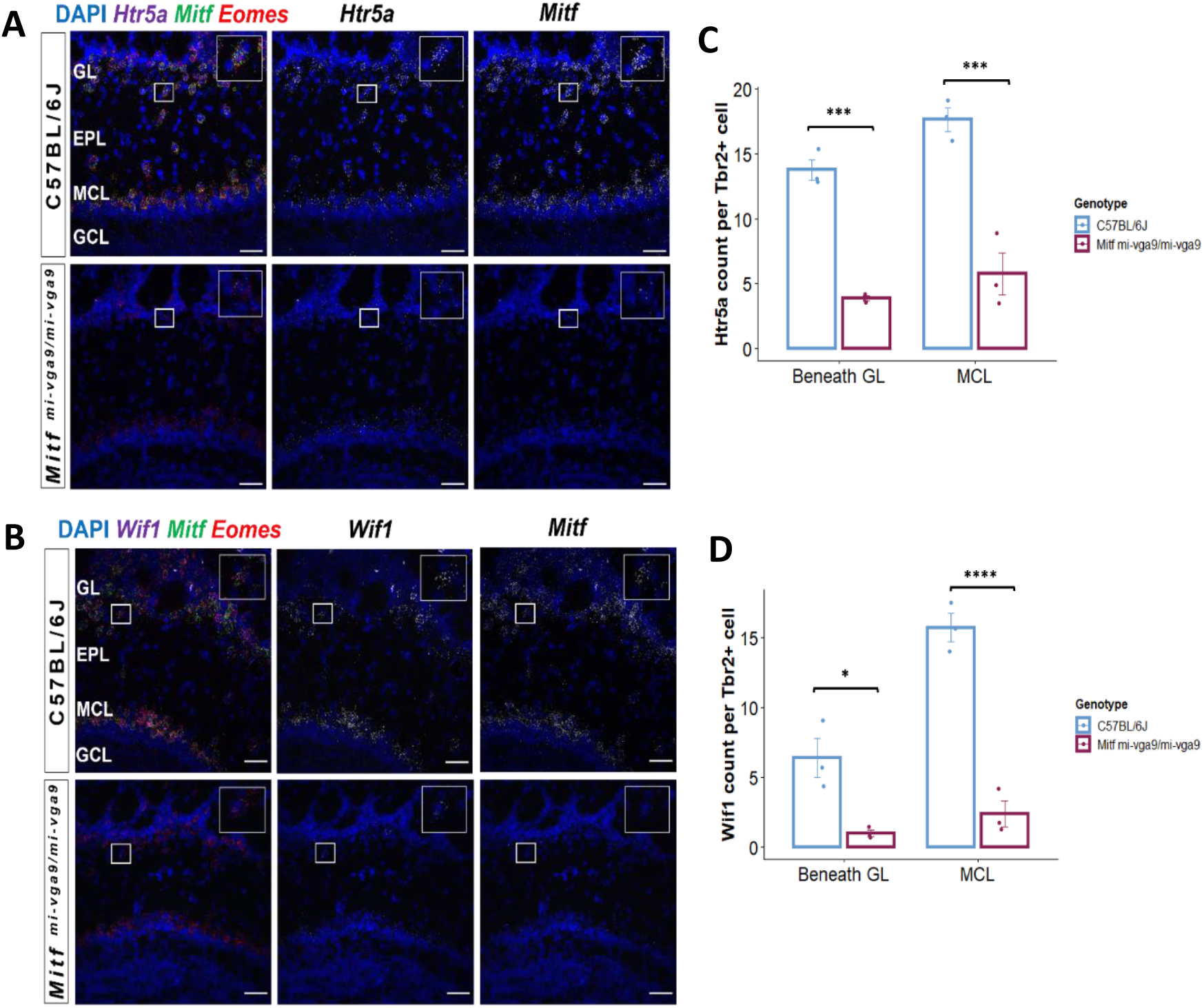
Changes in *Htr5a* and *Wif1* expression in *Mitf^mi-vga9/mi-vga9^* OBs. A, B. Representative smFISH images of *Htr5a* (A), and *Wif1* (B) expression in C57BL/6J and *Mitf^mi-vga9/mi-vga9^* OBs. mRNA transcripts of each gene of interest are shown in white. Scale bars indicate 50 μm. C, D. Quantification of *Htr5a* (C), and *Wif1* (D) mRNA transcripts beneath the GL and in the MCL of C57BL/6J and *Mitf^mi-vga9/mi-vga9^*OBs. N=3 per genotype. The values on the graphs indicate mean ± SE. P-values were calculated using two-way ANOVA and adjusted with Sidak multiple comparison. *p<0.05, ***p<0.001, ****p< 0.0001. GL (glomerular layer), EPL (external plexiform layer), MCL (mitral cell layer), GCL (granule cell layer).

**Figure 3.**
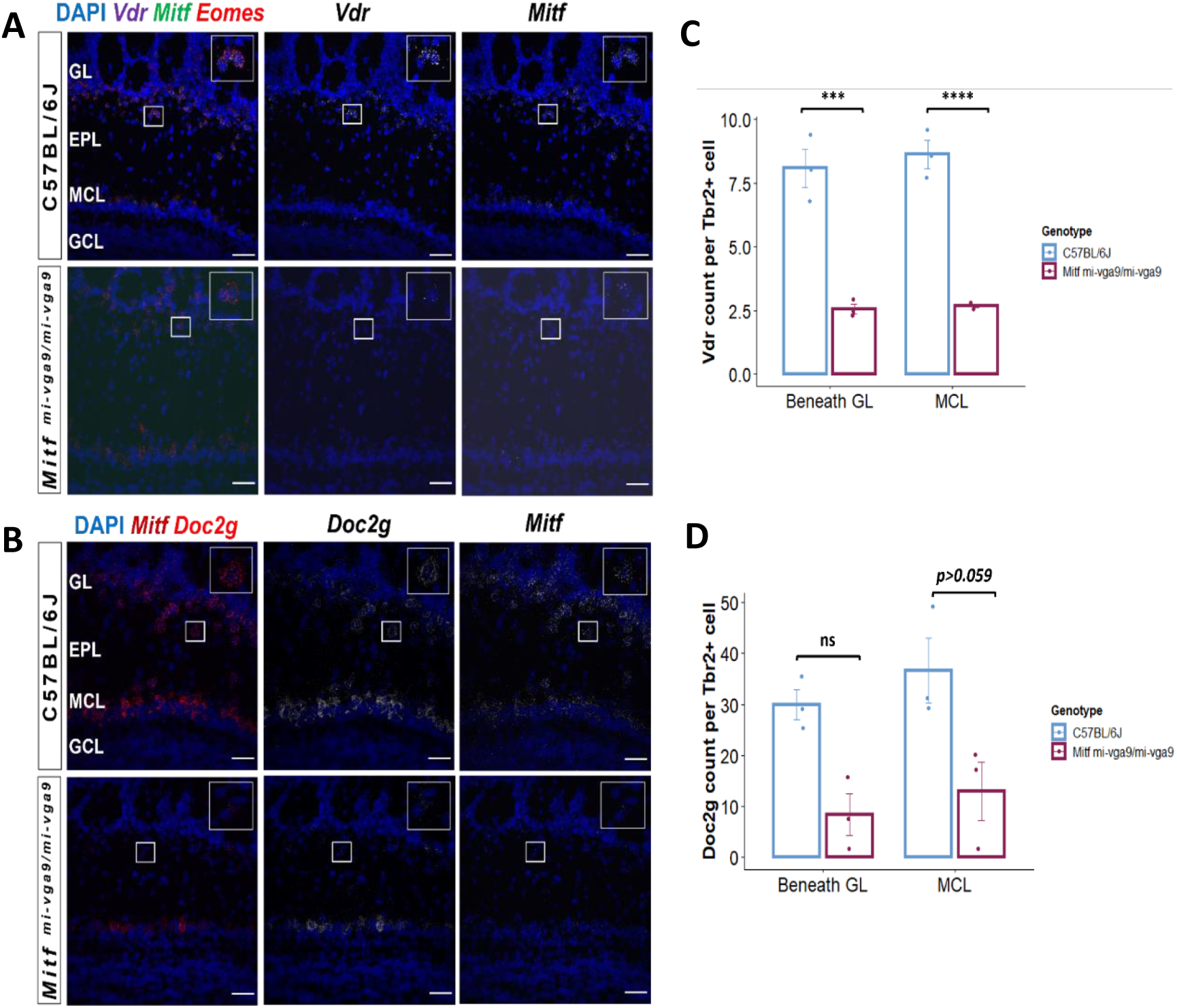
Changes in *Vdr* and *Doc2g* expression in *Mitf^mi-vga9/mi-vga9^* OBs. A, B. Representative smFISH images of *Vdr* (A), and *Doc2g* (B) expression in C57BL/6J and *Mitf^mi-vga9/mi-vga9^* OBs. mRNA transcripts of each gene of interest are shown in white. Scale bars indicate 50 μm. C, D. Quantification of *Vdr* (C), and *Doc2g* (D) mRNA transcripts beneath the GL and in the MCL of C57BL/6J and *Mitf^mi-vga9/mi-vga9^* OBs. N=3 per genotype. The values on the graphs are mean ± SE. P-values were calculated using two-way ANOVA and adjusted with Sidak multiple comparison. ***p<0.001, ****p<0.0001. GL (glomerular layer), EPL (external plexiform layer), MCL (mitral cell layer), GCL (granule cell layer); ns (not significant).

**Table 2.**
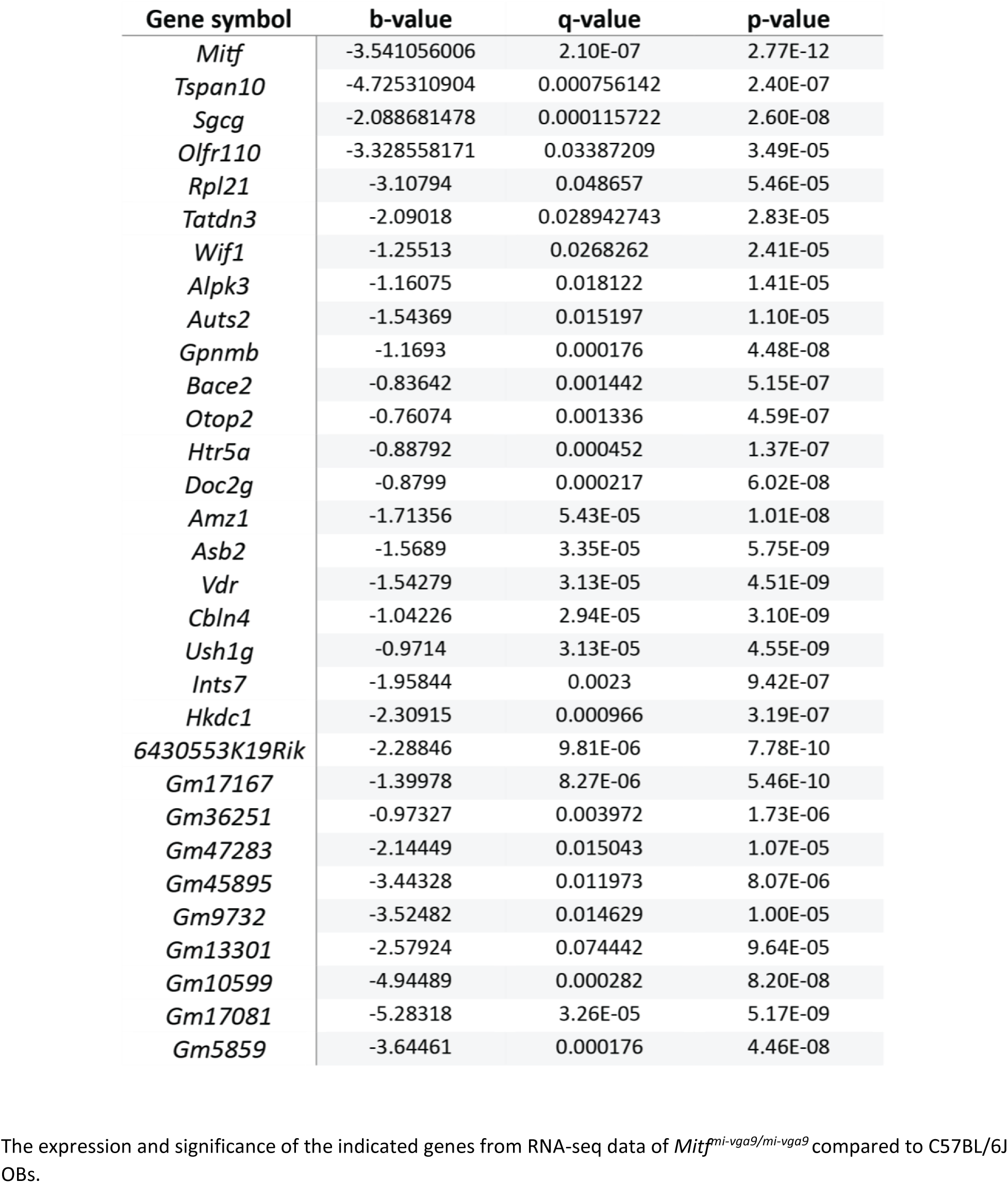
Genes with decreased expression in *Mitf^mi-vga9/mi-vga9^* OBs.

### *Mitf* affects the number of middle tufted cells and their gene expression

Using single-nucleus RNA sequencing (snRNA-seq), Zeppilli et al. (2021) identified distinct subgroups of mitral (M1-M3) and middle/external tufted (T1, ET1-ET4) cells based on gene expression (Figure 4A-D). In the present study, reduced expression of four middle tufted cells (mTCs; T1 subgroup) defining genes (*Sgcg*, *Tspan10*, *Vdr*, *Olfr110*) was observed in the OBs of *Mitf^mi-vga9/mi-vga9^* mice compared to C57BL/6J controls. This reduction was statistically significant for each gene: *Tspan10* (Figure 4E), *Vdr* (Figure 4F), *Sgcg* (Figure 4G), and *Olfr110* (Figure 4H). As expression levels were generally low, smFISH was used to determine expression at the cellular level. In *Mitf^mi-vag9/mi-vga9^* OBs, *Sgcg* expression decreased 88% beneath the GL and 93% in the EPL of *Mitf^mi-vga9/mi-vga9^* OBs as compared to C57BL/6J OBs (Figure 5A, B; *Sgcg:* beneath GL: *t _(24)_ = 9.908, p = 0.0001*; EPL: *t _(24)_ = 15.093, p = 0.0001,* Sidak multiple comparison). *Tspan10* expression was also decreased and was 83% less beneath the GL and 97% less in the EPL of *Mitf^mi-vga9/mi-vga9^* OBs (Figure 6A, B; *Tspan10:* beneath GL: *t _(24)_ = 5.878, p = 0.0001*; EPL: *t _(24)_ = 9.404, p = 0.0001,* Sidak multiple comparison).

**Figure 4.**
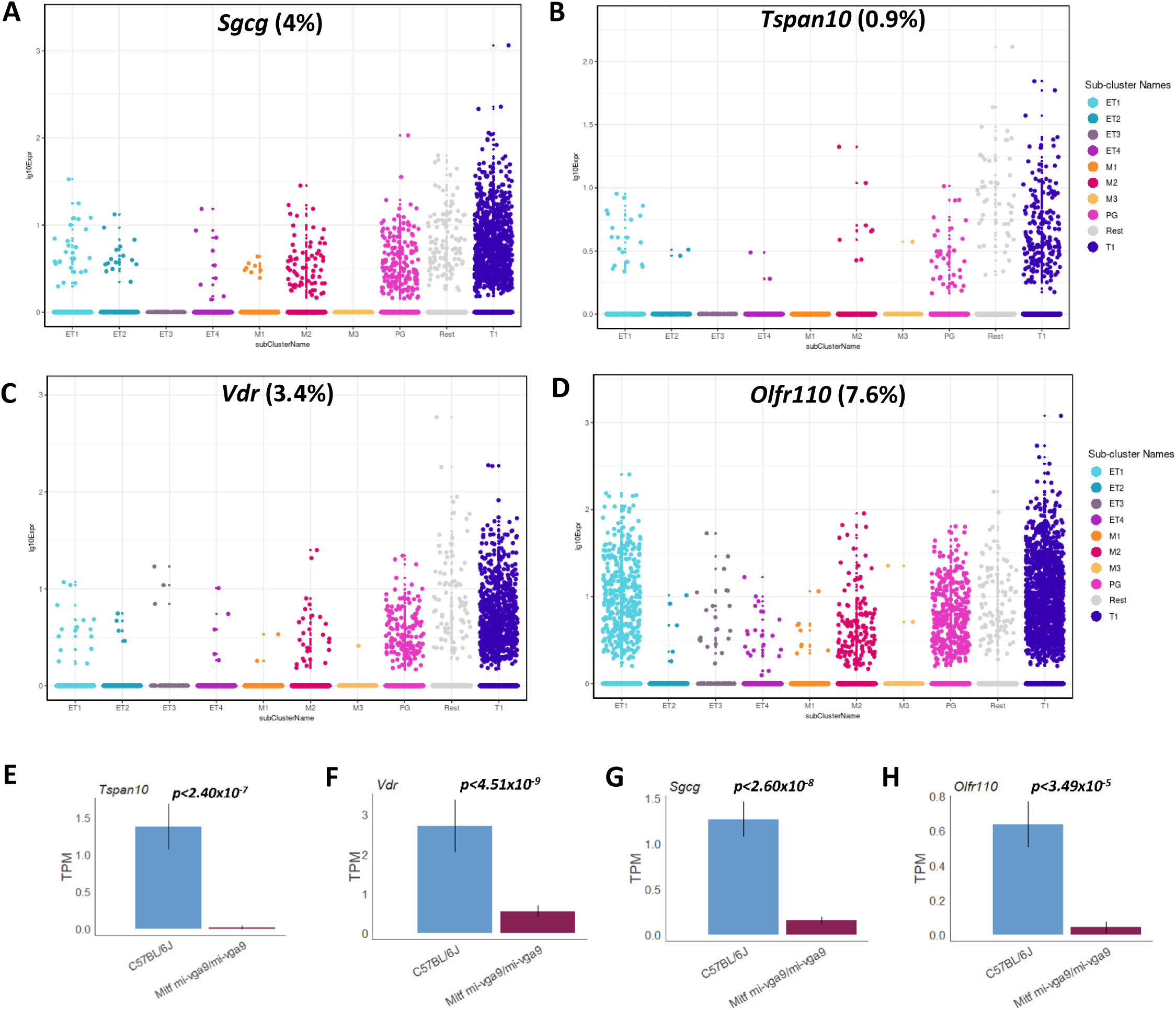
Reduced expression of mTCs genes (T1-subgroup) in *Mitf^mi-vga9/mi-vga9^* OBs. A-D. Dot plots showing expression of *Sgcg* (A), *Tspan10* (B), *Vdr* (C), *Olfr110* (D) in the different subgroups of PNs and in other cell types of the OB (Zeppilli et al., 2021). The percentage of cells expressing the gene of interest in the mTCs (T1 subgroup) is shown. E-H. Expression of the mTCs (T1)-specific genes *Tspan10* (E), *Vdr* (F), *Sgcg* (G), and *Olfr110* (H) in TPM in C57BL/6J and *Mitf^mi-vga9/mi-vga9^* OBs. N=4 per genotype. P-values on the graphs indicate the significance in *Mitf^mi-vga9/mi-vga9^* OBs compared to C57BL/6J OBs. ET1, ET2, ET3, ET4 (external tufted cell types 1-4); M1, M2, M3 (mitral cell types 1-3); PG (periglomerular cells); T1 (mTCs); TPM (transcript per million).

**Figure 5.**
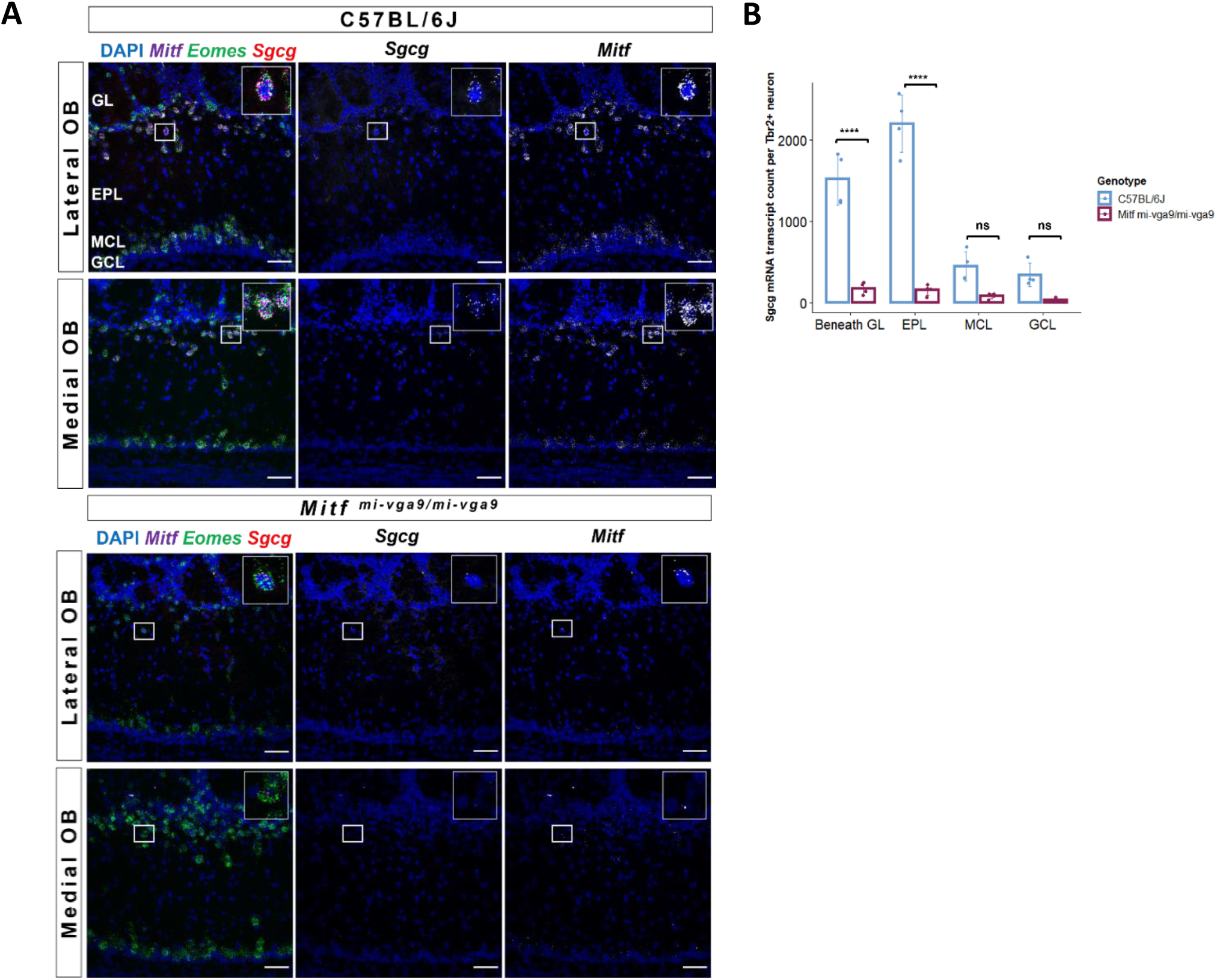
Reduced expression of *Sgcg* in mTCs of *Mitf^mi-vga9/mi-vga9^* OBs. A. Representative smFISH images of *Sgcg* in C57BL/6J and *Mitf^mi-vga9/mi-vga9^* OBs. mRNA transcripts of each gene are shown in white. B. Overall (lateral and medial OB) quantification of *Sgcg* mRNA transcripts beneath the GL, in the EPL, MCL and GCL of C57BL/6J and *Mitf^mi-vga9/mi-vga9^* OBs. The values on the graphs are mean ± SE. P-values were calculated using two-way ANOVA and adjusted with Sidak multiple comparison. N=4 per genotype. Scale bars indicate 50 μm. ****p<0.0001. GL (glomerular layer), EPL (external plexiform layer), MCL (mitral cell layer), GCL (granule cell layer); ns (not significant).

**Figure 6.**
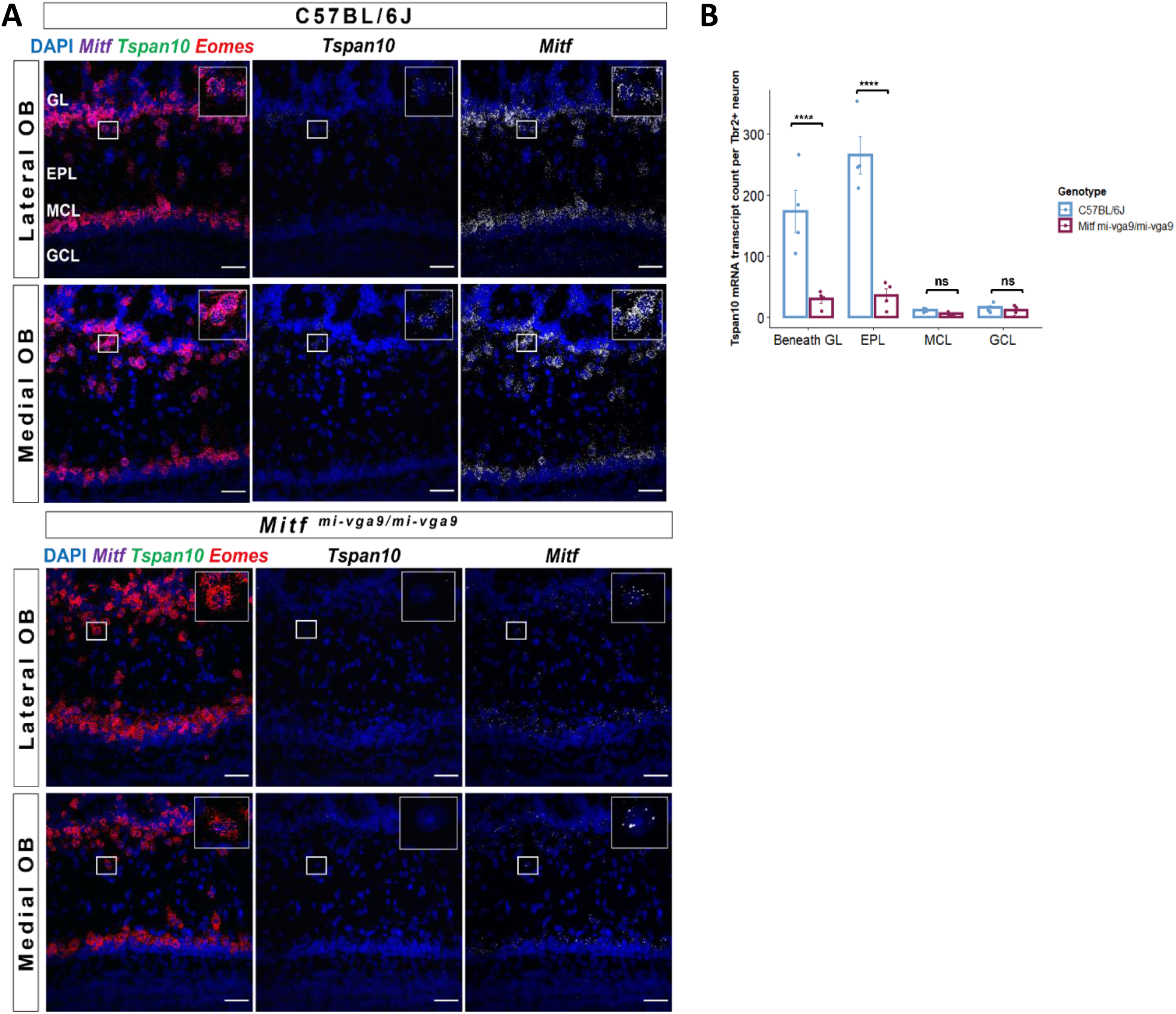
Expression of *Tspan10* is decreased in mTCs of *Mitf^mi-vga9/mi-vga9^* OBs. A. Representative smFISH images of *Tspan10* in C57BL/6J and *Mitf^mi-vga9/mi-vga9^* OBs. mRNA transcripts of each gene are shown in white. B. Overall quantification (lateral and medial OB) of *Tspan10* mRNA transcripts beneath the GL, in the EPL, MCL and GCL of C57BL/6J and *Mitf^mi-vga9/mi-vga9^* OBs. The values on the graphs are mean ± SE. P-values were calculated using two-way ANOVA and adjusted with Sidak multiple comparison. N=4 per genotype. Scale bars indicate 50 μm. ****p<0.0001. GL (glomerular layer), EPL (external plexiform layer), MCL (mitral cell layer), GCL (granule cell layer); ns (not significant).

Importantly, the expression of other mTCs marker genes (*Fam84b*, *Barhl2*, *Olfr111*, *Cacna1g*, *Kcna10*) was not significantly reduced in the sequencing data, indicating the presence of this neuronal population in *Mitf^mi-vga9/mi-vga9^* OBs. However, analysis of the ABA showed that most genes suggested to be specific to mTCs, have a broader expression than observed in the study of Zeppilli and associates (2021). In fact, only *Fam84b* showed specific expression in mTCs in the ABA database. To ascertain the presence of mTCs, smFISH for *Fam84b* was therefore performed on *Mitf^mi-vga9/mi-vga9^*OBs. Consistent with the ABA and RNA-seq data, *Fam84b* is expressed specifically in *Tbr2/Eomes* positive cells in the EPL (Figure 7A) and its expression is decreased by 70% in *Mitf^mi-vga9/mi-vga9^* OBs compared to C57BL/6J OBs (Figure 7B; *Fam84b:* EPL: *t _(24)_ = 6.161, p = 0.0001*). This demonstrates that not all specific mTCs marker genes are absent in the *Mitf^mi-vga9/mi-vga9^*OBs and therefore confirms the presence of mTCs in *Mitf^mi-vga9/mi-vga9^*OBs.

**Figure 7.**
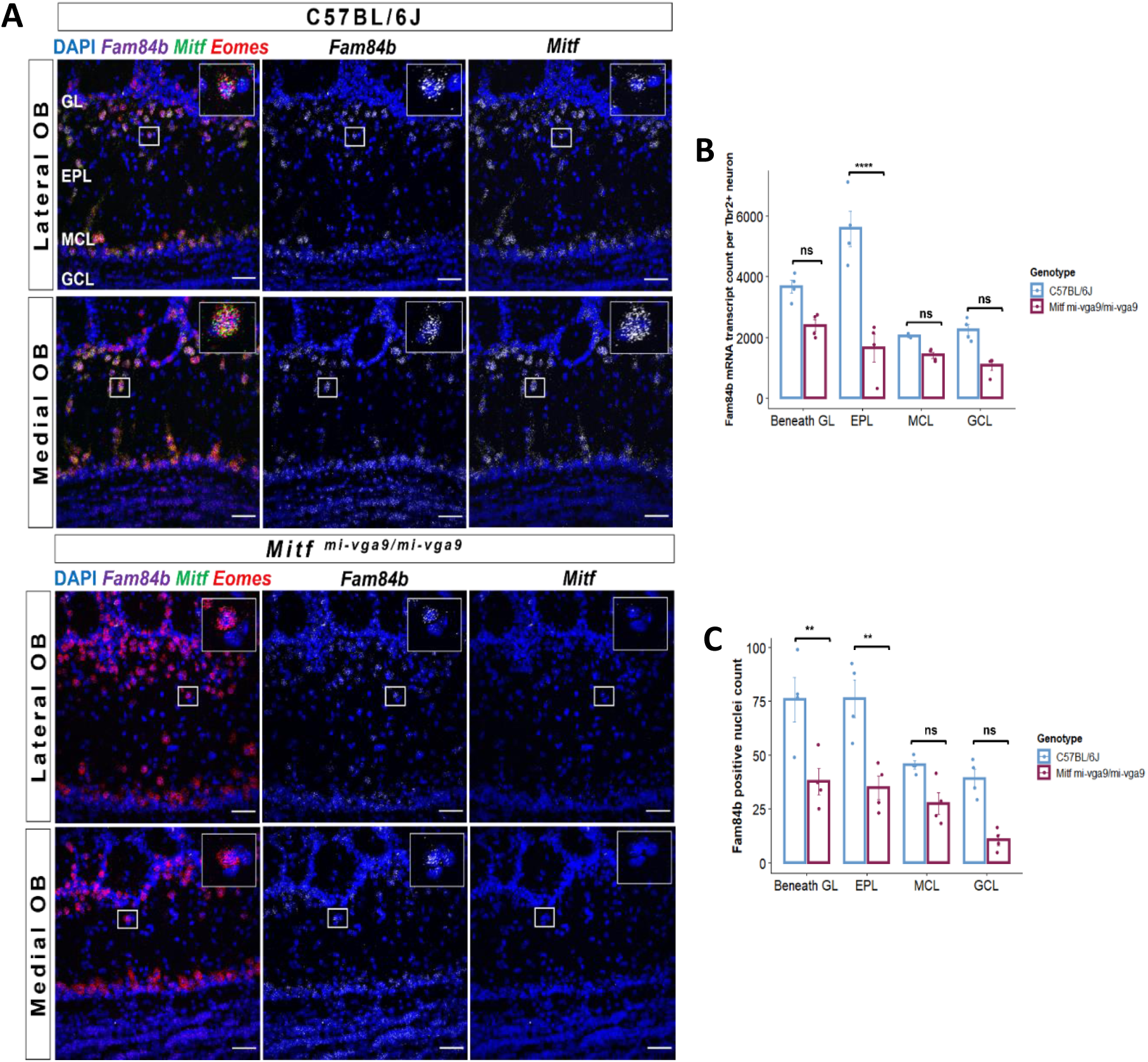
MTCs are present but reduced in number in *Mitf^mi-vga9/mi-vga9^* OBs. A. Representative smFISH images of *Fam84b* in C57BL/6J and *Mitf^mi-vga9/mi-vga9^* OBs. mRNA transcripts are shown in white. B. Quantification of *Fam84b* mRNA transcripts beneath the GL, in the EPL, MCL and GCL of C57BL/6J and *Mitf^mi-vga9/mi-vga9^* OB. N=4 per genotype. C. Quantification of *Fam84b* positive cells (clusters) beneath the GL, in the EPL, MCL and GCL of C57BL/6J and *Mitf^mi-vga9/mi-vga9^* OBs. N=4 per genotype. The values on the graphs are mean ± SE. P-values were calculated using two-way ANOVA and adjusted with Sidak multiple comparison. Scale bars indicate 50 μm. **p<0.01, ****p<0.0001. GL (glomerular layer), EPL (external plexiform layer), MCL (mitral cell layer), GCL (granule cell layer); ns (not significant).

*Fam84b* positive cells were also counted and showed a significant decrease in the beneath the GL (50%) and the EPL (54%) of *Mitf^mi-vga9/mi-vga9^*OBs compared to C57BL/6J OBs, whereas the decrease was not significant in the MCL or GCL where mTCs are not located (Figure 7C; beneath GL: *t _(24)_ = 4.177*, *p = 0.0094*; EPL: *t _(24)_ = 4.205*, *p = 0.0087*, Sidak multiple comparison).

Taken *together, Fam84b* positive mTCs are still present in *Mitf^mi-vga9/mi-vga9^* OBs, their numbers are reduced and the expression of *Fam84b* is decreased in individual cells. The near absence of *Sgcg* and *Tspan10* expression from the *Mitf^mi-vga9/mi-vga9^* OBs suggests that they are candidate MITF target genes in mTCs. MITF binding is found in both genes in melanoma cells, supporting them as targets of MITF in neurons (Supplementary figure 1B, C).

### MITF overexpression leads to increased expression of *Tspan10*

To examine the direct effect of MITF on the expression of the putative target genes, MITF was overexpressed in the N2a neuronal cell line (Figure 8A). RNA-seq analysis was performed to evaluate gene expression changes in N2a cells expressing MITF compared to cells expressing empty vector (EV) (Figure 8B). This analysis showed that MITF expression affected well-characterized melanocyte specific MITF target genes, such as *Pmel* and *Trpm1* (Figure 8C). Importantly, MITF overexpression led to a manyfold increase in expression of *Tspan10* (Figure 8D). Earlier studies have found a connection between *Mitf* and *Tspan10* in melanoma (Rambow et al., 2015) and in osteoclasts (Zhou et al., 2014). This further supports the hypothesis that *Tspan10* is an MITF target gene in mTCs and other cell types.

**Figure 8.**
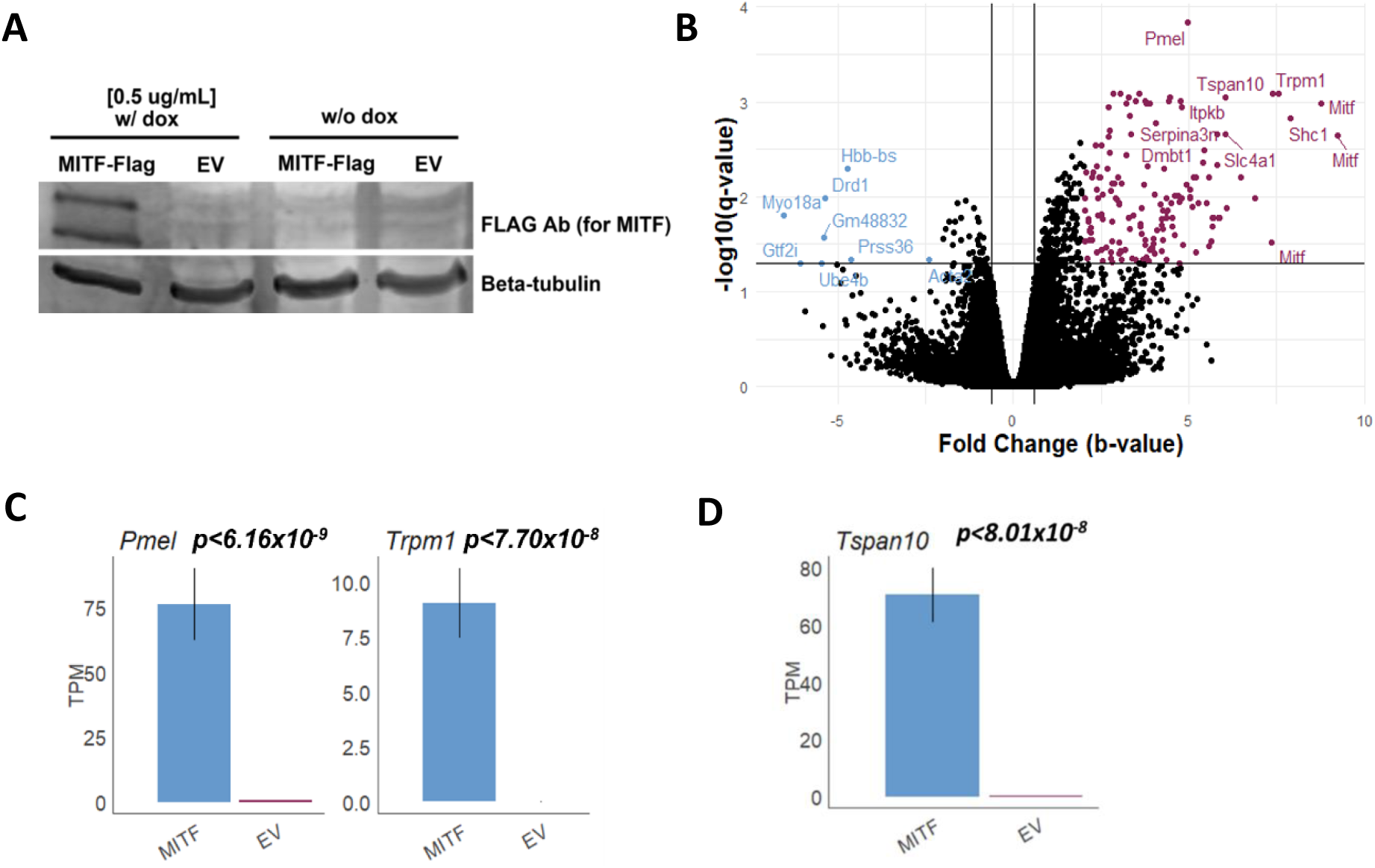
Expression of the *Tspan10* mRNA is increased in N2a cell line upon MITF overexpression. A. Representative western blot image showing MITF-Flag expression after inducing MITF overexpression with 0.5 μg/mL dox for 24 hours. Also is shown the same cells transfected with the MITF-Flag construct but not treated with dox. B. Volcano plot showing differentially expressed genes upon overexpressing MITF-Flag in the N2a cells. Comparison was made to cells expressing EV following dox induction. Blue represents decreased genes and purple represents increased genes. C. Expression of MITF target genes in melanocytes (*Pmel, Trpm1*) in N2a cell line expressing MITF-Flag or EV with dox, indicated in TPM. D. Expression of *Tspan10* RNA in N2a cell line expressing MITF-Flag or EV with dox, indicated in TPM. N=3 per condition. P-values (*p<0.05) indicate the significance in N2a cells expressing MITF-Flag compared to EV. Dox (doxycycline); TPM (transcript per million).

### Putative MITF target genes are not activity dependent

One model of the role of *Mitf* in M/T cells is that *Mitf* negatively affects neuronal activity via intrinsic mechanisms, contributing to homeostatic plasticity or alterations in gain/control (Atacho et al., 2020). Briefly, the model proposes that following neuronal activity, increased I_A_ currents mediated by MITF lead to reduction in neuronal activity of M/T cells, and thus adaptation to the odorant. In *Mitf^mi-vga9/mi-vga9^* mice olfactory adaptation is altered (Atacho et al., 2020) similar to what has been shown in other mouse mutants in which M/T cells are hyperactive (Fadool et al., 2004).

As the loss of *Mitf* leads to hyperactivity in primary M/T cells (Atacho et al., 2020), it is also possible that the genes affected in the current study are reduced in expression due to the changes in neuronal activity. Alterations in gene expression were examined using RNA-seq comparing C57BL/6J and *Mitf^mi-vga9/mi-vga9^*OBs, following exposure to amyl acetate (AA), an odorant that activates multiple glomeruli (Bepari et al., 2012; Rubin & Katz, 1999; Wu et al., 2017). Two experimental setups were used to determine whether the expression of the putative MITF target genes would change upon activity induction. In the first setup, mice were exposed to AA for 30 minutes, while in the second setup olfactory exposure was followed with a 30-minute rest to target different transcriptional events in the OB response (Figure 9A). For gene expression analysis, odor-exposed mice of each genotype were normalized with their corresponding control group.

**Figure 9.**
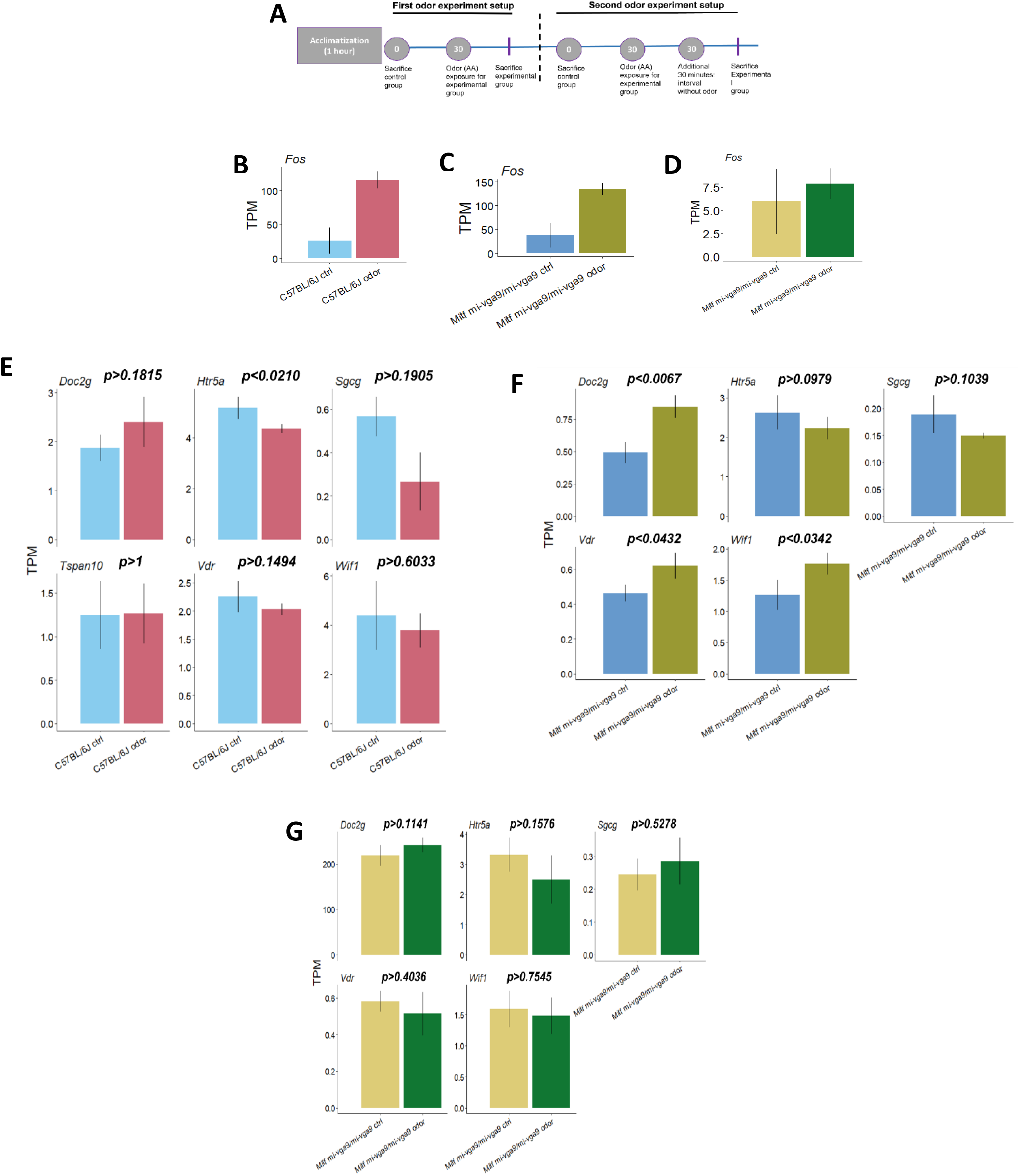
Putative MITF target genes are not activity dependent. A. Schematic representation of an odor-exposure experiments. B-D. The expression of *c-fos (Fos)* after 30 minutes odor exposure without a time interval in C57BL/6J (B) and *Mitf^mi-vga9/mi-vga9^* (C) OBs and with a time interval in *Mitf^mi-vga9/mi-vga9^* OBs (D). N= 3 per genotype. E. The expression of putative MITF target genes (in TPM) after 30 minutes of odor exposure without a time interval in C57BL/6J OBs. N= 3 per condition. F. The expression of putative MITF target genes (in TPM) after 30 minutes of odor exposure without a time interval in *Mitf^mi-vga9/mi-vga9^* OBs. N= 3 per condition. G. The expression of putative MITF target genes in TPM after 30 minutes of odor exposure with a time interval in *Mitf^mi-vga9/mi-vga9^* OBs. N=3 per condition. P-values (*p<0.05) on the bar graphs indicate the significance in C57BL7/6J odor compared to C57BL/6J control and *Mitf^mi-vga9/mi-vga9^* odor compared to *Mitf^mi-vga9/mi-vga9^* control. TPM (transcript per million).

The well-characterized neuronal activity marker *c-fos* (Morgan & Curran, 1988) showed the expected expression increase in C57BL/6J and *Mitf^mi-vga9/mi-vga9^*OBs after odor exposure (Figure 9B-D), confirming odor-evoked neuronal activity. Subsequently, the expression of the putative MITF target genes was examined, comparing the experimental group of C57BL/6J and *Mitf^mi-vga9/mi-vga9^* mice to their respective control groups. *Htr5a* decreased slightly in C57BL/6J OBs (Figure 9E) while *Doc2g, Vdr*, and *Wif1* showed increased expression in *Mitf^mi-vga9/mi-vga9^* OBs (Figure 9F) after odor exposure. Despite the increase after odor expression, the changes in TPMs between odor exposed and control groups are minimal, thus suggesting limited biological significance. Their expression change was not comparable to changes in *c-fos* expression following odor exposure (Figure 9B-D). The analysis of gene expression at 30 and 60 minutes, revealed no changes in expression of putative target genes between C57BL/6J (data not shown) and *Mitf^mi-vga9/mi-vga9^* OBs (Figure 9G). Overall, none of the putative MITF target genes appear to be activity dependent, at the time scale examined.

## Discussion

We have identified several putative MITF target genes in PNs of the mouse OB based on their reduced expression in *Mitf^mi-vga9/mi-vga9^*mice, specific expression in PNs, and the presence of MITF binding peaks in non-neuronal cells. Further experiments are required to confirm actual binding in OB neurons and direct regulation of these genes by MITF. The putative MITF target genes in the mouse OB fall into two groups: those expressed widely in M/T cells (*Htr5a*, *Doc2g*, *Wif1*) and genes expressed selectively in subgroup of tufted cells (*Sgcg*, *Tspan10*, *Vdr*, *Olfr110*). The genes expressed broadly in M/T cells all appear to have a potentially negative impact on neuronal activity. The activation of *Htr5a* decreases neuronal excitability and neurotransmission which contributes to the overall modulation of neuronal activity (Guan et al., 2016; Kinsey et al., 2001; Yu et al., 2012). Similarly, *Wif1* inhibits the stimulatory effects of the Wnt pathway, modulating neuronal activity (Diep et al., 2004; Inestrosa & Toledo, 2008). The function of *Doc2g* has not yet been characterized, but it likely plays a role in neurotransmitter release similar to its homologs *Doc2*a and *Doc2b* (Fukuda & Mikoshiba, 2000; Sudhof, 2004). Of these genes only *Htr5a* has MITF binding peaks in ChIP-seq and CUT&RUN data from melanoma cells (Supplementary figure 1A; Laurette et al. 2015; Dilshat et al. 2021; Strub et al., 2011), thereby supporting it as a potential MITF target gene. The reduction in expression of these genes may in part explain the hyperactivity of primary cells from *Mitf^mi-vga9/mi-vga9^* OB. Hence, the role of *Mitf* in affecting neuronal activity of OB PNs appears to be complex, and to involve both synaptic and intrinsic mechanisms. This effect does not appear to be activity dependent as following olfactory activity, global gene expression analysis did not show any clear changes.

The expression of the second group of genes affected by *Mitf* (*Sgcg, Tspan10, Olfr110, Vdr*) is specific to a subgroup of tufted cells (middle tufted cells - mTCs) and have been identified as marker genes for these neurons in the OB (Zeppilli et al., 2021). The number of mTCs is reduced in *Mitf^mi-vga9/mi-vga9^*OBs. This suggests that MITF plays a role in the function of mTCs or their development. SmFISH *shows* that *Sgcg, Tspan10* and *Vdr* are expressed distinctly in mTCs in C57BL/6J OBs. *Sgcg* and *Tspan10* are nearly absent and *Vdr* significantly decreased in *Mitf^mi-vga9/mi-vga9^* OBs. *Vdr* is required for the activity inducing effects of 1,25(OH)2D in the cortex (Gooch et al., 2019) and thus might play a specific role in mTCs of the OB. The olfactory receptor gene *Olfr110* is expressed outside the olfactory epithelium and appears to play functional roles in neurons (Gaudel et al., 2019). As *Vdr* and *Olfr110* do not have MITF binding peaks in non-neuronal cells, changes in their expression might be secondary effects.

*Sgcg*, encodes the SGCG protein which is primarily found in striated muscle (Ervasti & Campbell, 1991; Yamamoto et al., 1993) where it forms heterotetrameric complexes with other sarcoglycans. It is crucial for assembling the dystrophin-associated glycoprotein (Ozawa et al., 1995). Mutations in *Sgcg* are linked to cardiomyopathy and muscular dystrophy in humans (Noguchi et al., 1995). MITF binding peaks are found in the *Sgcg* in melanoma cells (Supplementary figure 1B; Laurette et al. 2015; Dilshat et al. 2021; Strub et al., 2011). While the role of SGCG in neurons remains unclear, it is likely to play a role in cell-cell interactions.

The strongest candidate for an MITF target gene in mTCs is *Tspan10*. Overexpression of MITF in N2a cells increases the expression of *Tspan10*; MITF peaks are found in the gene in melanoma cells (Supplementary figure 1C; Laurette et al. 2015; Dilshat et al. 2021; Strub et al., 2011). It has also been associated with a regulatory network related to MITF-mediated migration of melanoma cells (Seong et al., 2012). The best-known role of *Tspan10* is activating ADAM10 metalloprotease during osteoclastogenesis (Zhou et al., 2014). Although the role of ADAM10 is not fully understood in the CNS, studies have shown its involvement in the regulation of neurogenesis, embryonic axonal growth, and at the synapses (Kuhn et al., 2016; Malinverno et al., 2010; Musardo et al., 2014). Moreover, ADAM10 is important for melanocyte attachment in the epidermis and its overexpression drives melanoma progression (Lee et al., 2010; Tharmarajah et al., 2012). Rambow et al. (2015) showed that *Tspan10* expression is increased in multiple melanoma cell lines. It is plausible that the role of *Tspan10* in mTCs relates to the specific function of these cells in the olfactory circuitry and could involve regulation of cell-cell interactions.

The SGCG, TSPAN10, and OLF110 proteins are membrane proteins that might be involved in cell-cell communication processes such as axonal pathfinding or wiring of the mTCs. If this is the case, alterations in the connectivity between the OBs and higher olfactory regions are to be expected. In the olfactory cortex (OCx) mTCs extend axons to the olfactory nucleus (AON) and the olfactory tubercle (OT) while mitral cell innervation varies across AON, OT, the anterior and posterior piriform cortex, the medial amygdaloid nucleus and the anterior and posterior cortical amygdaloid nuclei (Imamura et al., 2020). Further analysis of the link between OB and OCx, along with analysing conditional knock-out mice is likely to clarify the distinct role of MITF for the functional specificity of these neurons.

The MITF transcription factor is central for the development and function of melanocytes, mast cells and osteoclasts. It regulates multiple pathways such as differentiation, proliferation, and apoptosis through the expression of genes specific to each cell type. This study highlights the diverse effects of the loss of *Mitf* on gene expression in the OB where it plays at least two roles: regulating expression of genes which affect neuronal activity in M/T cells and genes which define and likely play specific roles in a subpopulation of tufted cells, the mTCs.

## Acknowledgements

This study was supported by Research Fund of Iceland to PHP (217945-052) and a grant from the University of Iceland Doctoral Grants Fund to FM. We thank the Biomedical Center at the University of Iceland (BMC). We are also grateful to our colleagues from the BMC, especially Ragnhildur Káradóttir, Margrét Helga Ögmundsdóttir, Sigríður Rut Franzdóttir, Sana Gadiwalla and Nhung Hong Vu for their contribution to the project.

---

**Supplementary figure 1.**
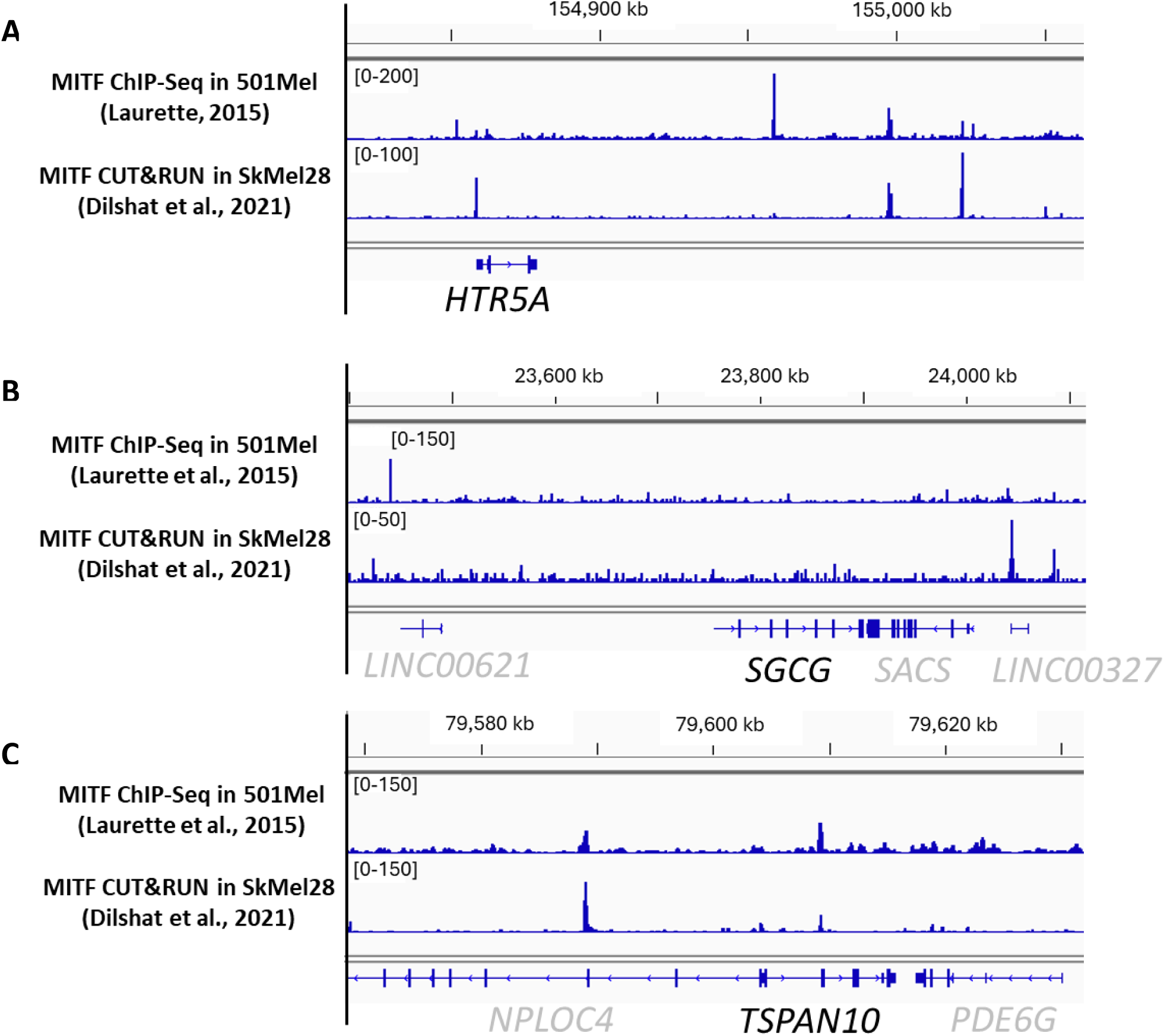
MITF peaks in the *HTR5A, SGCG,* and *TSPAN10* genes in melanoma cells. A-C. IGV Tool images showing MITF binding peaks on the human *HTR5A* (A), *SGCG* (B) and *TSPAN10* (C) genes based on MITF ChIP-seq data from Laurette et al., (2015) and CUT&RUN data from Dilshat et al., (2021).

